# Computational Design of Asymmetric Three-dimensional RNA Structures and Machines

**DOI:** 10.1101/223479

**Authors:** Joseph D. Yesselman, Daniel Eiler, Erik D. Carlson, Alexandra N. Ooms, Wipapat Kladwang, Xuesong Shi, David A. Costantino, Daniel Herschlag, Michael C. Jewett, Jeffrey S. Kieft, Rhiju Das

**Author notes:** Correspondence should be addressed to R.D.

## Abstract

The emerging field of RNA nanotechnology seeks to create nanoscale 3D machines by repurposing natural RNA modules, but successes have been limited to symmetric assemblies of single repeating motifs. We present RNAMake, a suite that automates design of RNA molecules with complex 3D folds. We first challenged RNAMake with the paradigmatic problem of aligning a tetraloop and sequence-distal receptor, previously only solved via symmetry. Single-nucleotide-resolution chemical mapping, native gel electrophoresis, and solution x-ray scattering confirmed that 11 of the 16 ‘miniTTR’ designs successfully achieved clothespin-like folds. A 2.55 Å diffraction-resolution crystal structure of one design verified formation of the target asymmetric nanostructure, with large sections achieving near-atomic accuracy (< 2.0 Å). Finally, RNAMake designed asymmetric segments to tether the 16S and 23S rRNAs together into a synthetic singlestranded ribosome that remains uncleaved by ribonucleases and supports life in *Escherichia coli*, a challenge previously requiring several rounds of trial-and-error.

## Introduction

RNA-based nanotechnology is an emerging field that harnesses RNA’s unique structural properties. Perhaps more so than for other biomolecules, RNA’s tertiary structure is composed of discrete and recurring components known as tertiary ‘motifs’ (1–4). Along with the helices that they interconnect, many of these structural motifs appear highly modular; that is, each motif folds into a common, well-defined 3D structure in a large range of contexts (5–7). By exploiting symmetry, motif repetition, and expert modeling, these motifs have been assembled into rationally designed structures, including polygons, cubes, and sheets (8–14). Nevertheless, the RNA molecules generated to date are mostly symmetrical and structurally simple, lacking the complexity observed in natural RNA machines, which contain a larger repertoire of distinct interacting motifs. The importance of this structural diversity is implied by the wide range of functionality observed in naturally occurring RNAs (15–18). Rationally designing RNAs with increased functional diversity will require methods to generate asymmetric folds with a larger set of motifs (19).

Including additional structural motifs in rational 3D RNA design is challenging for two reasons. First, introducing new motifs demands rigorous testing for modular behavior, i. e., determining if the motifs maintain their 3D folds when installed into new RNAs. Second, with current methods and software, the rational 3D design of RNA can be laborious, even with a limited set of motifs, as the designer is required to intuit the correct helical separation between motifs, ensure that tertiary contacts are in the proper orientation, and guarantee the correct formation of secondary structure.

Here, we address these issues by introducing a method to design continuous chains of RNA motifs that twist and translate between any two desired helical endpoints that can harbor functional elements or tertiary contacts (Figure 1). We then demonstrate two proofs of concept using design problems that have been challenging in prior studies. First, to allow for stringent structural tests, we tackle the classic problem of designing an RNA that aligns the two parts of the tetraloop/tetraloop-receptor (TTR) so that they can form the TTR tertiary contact upon RNA chain folding (Figure 1b). This 3D design challenge was the first problem solved in RNA nanotechnology (7), but all solutions to date have either relied on symmetric design principles to guide dimer, fiber, or higher-order assembly (20) or have resulted in a tertiary structure with minimal stability (21). In addition, these prior solutions required manual modeling by experts. Second, we demonstrate that computational 3D design can aid in difficult alignment challenges encountered in the development of functional RNA machines (22–24). For example, design challenges involving ribosomal subunit tethering (22, 23) previously proved difficult to address using manual modeling, as they require complex RNA segments to correctly orient RNA nanostructures relative to one another in 3D space. To demonstrate the broad application of computational 3D design, we build and test novel RNA segments that tether and align the small and large subunits of the ribosome together into a single RNA strand *in vivo*; this design problem previously required more than a year to solve using manual design and trial-and-error refinement based on *in vivo* assays (25).

**Figure 1:**
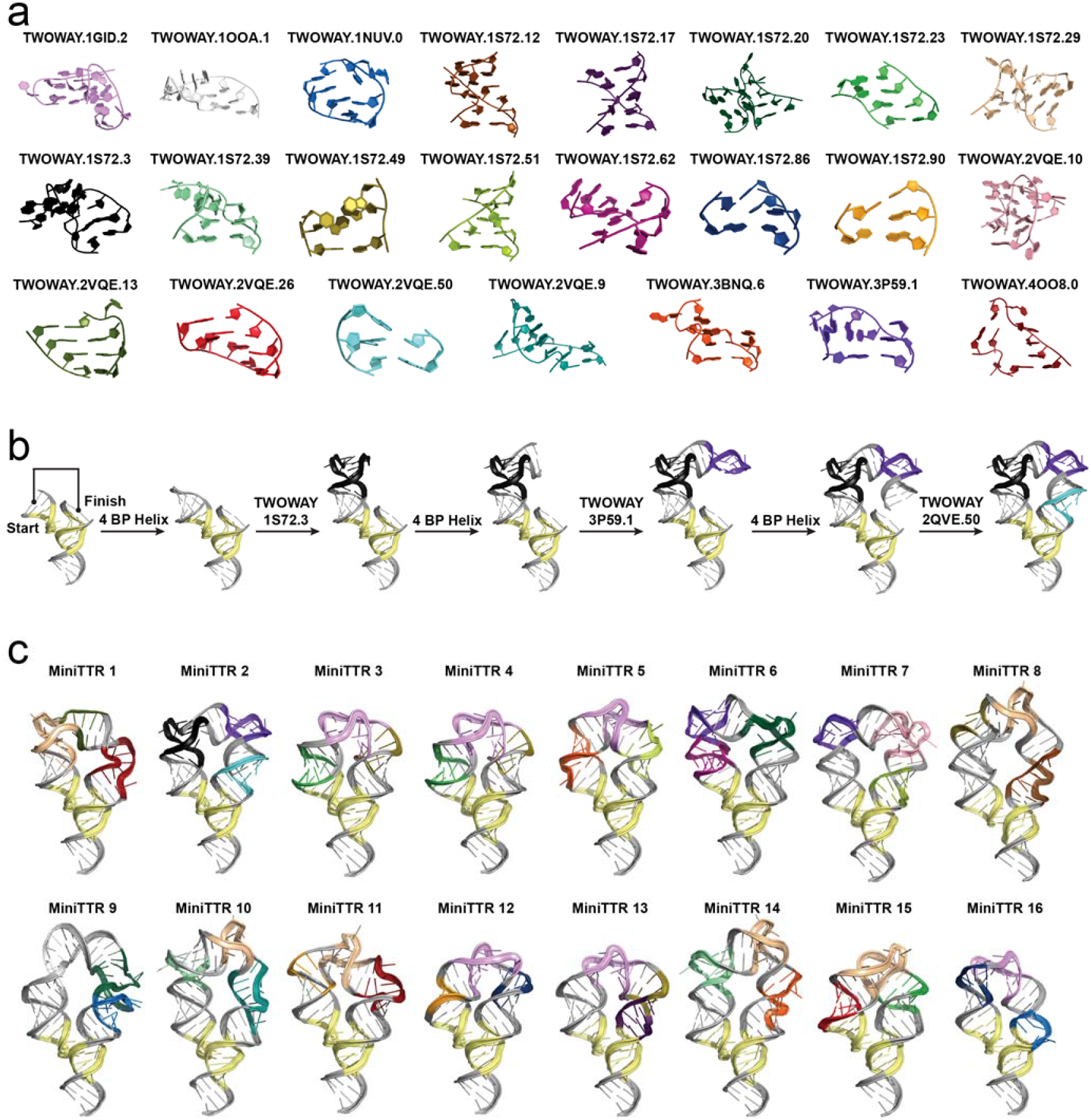
a) All unique motifs included in the miniTTR designs (See SI Table 2 for common names where available). b) Demonstration of RNAMake’s design algorithm, which builds an RNA path via the successive addition of motifs and helices from the starting base pair to the ending base pair. c) Ensemble of all RNAMake miniTTR designs. Each is colored according to the motifs (panel (a)) that were used to generate it.

Both the new minimal TTR (miniTTR) and ribosome tether constructs were generated using a fully automated toolkit for motif-based design that we call RNAMake. This toolkit contains a library of hundreds of unique motifs and a path-finding algorithm to connect any two points in 3D space at a desired relative orientation using an RNA segment. Our tests demonstrate that purely computational design can make use of multiple structurally diverse motifs within a single strand of RNAs to generate asymmetric, monomeric structures that fold correctly *in vitro* and to reengineer machines that support life.

Furthermore, our results confirm the modularity of dozens of novel non-canonical motifs not previously utilized in design.

## Results

### Generation and Use of a Curated Structural Motif Library

To expand the library of available motifs for rational design, we collected motifs from all unique RNA structures that had been deposited in a crystallographic database (see Methods). The final set contained two-way junctions, higher-order junctions, hairpins, and tertiary contacts with 461, 61, 290, and 89 unique motifs respectively. To efficiently sample over the large set of motif modules in our library, we developed a path-finding search algorithm. This algorithm models canonical helical segments of 1 to 22 base pairs that interconnect noncanonical structural motifs (see Methods). The canonical helical segments are idealized and sequence invariant (26), but after the completion of 3D structural design, they are filled in with sequences that best match the target secondary structure and minimize alternative secondary structures. This optimization of the helical sequences is carried out with EteRNAbot, developed previously through internet-scale crowdsourcing and tested through chemical mapping (27).

### RNAMake Automates RNA 3D Design That Includes Diverse Motifs

The assessment of the structural accuracy of RNAMake’s design ability is the first step toward using it as a reliable tool to generate functional RNA machines. We assessed RNAMake’s ability to generate structurally accurate 3D RNA structures by challenging it to align the tetraloop and receptor parts of the TTR in their bound conformation using a designed RNA segment. To generate these miniTTR constructs, we first extracted the coordinates from the X-ray crystal structure of the TTR from the P4-P6 domain of the *Tetrahymena* ribozyme (residues 146-157, 221-246, and 228-252 from PDB 1GID) (28).

Second, we used RNAMake to build structural segments composed of two-way junctions and helices spanning the last base pair of the hairpin (A146-U157) to base pair U221-A252 of the tetraloop-receptor, connecting the TTR into a single continuous strand (Figure 1b). Of 200,000 RNA segments generated, sixteen were selected based on two criteria: 1) fewest motifs per solution (three unique tertiary motifs) and 2) tightest alignment of the two segments of the TTR to their target spatial and rotational orientations. These computational designs ranged from 75 to 102 nucleotides in size (for full sequences, see SI Table 1), significantly shorter than the 157 nucleotides of the natural P4-P6 domain RNA.

**Table 1:** Biochemical assay results for each miniTTR.

| Construct | Chemical Mapping | Mobility Assay | Mg<sup>2+</sup> Titration |
| --- | --- | --- | --- |
| miniTTR 1 | Success | Success | Success |
| miniTTR 2 | Success | Success | Success |
| miniTTR 3 | Partial | Failure | Failure |
| miniTTR 4 | Partial | Success | Failure |
| miniTTR 5 | Success | Success | Success |
| miniTTR 6 | Success | Success | Success |
| miniTTR 7 | Success | Success | Success |
| miniTTR 8 | Failure | Failure | Failure |
| miniTTR 9 | Success | Success | Success |
| miniTTR 10 | Success | Success | Success |
| miniTTR 11 | Success | Success | Success |
| miniTTR 12 | Success | Success | Success |
| miniTTR 13 | Failure | Failure | Failure |
| miniTTR 14 | Success | Success | Success |
| miniTTR 15 | Success | Multimer | Success |
| miniTTR 16 | Success | Success | Success |

The miniTTR designs included 23 unique motifs (Figure 1a and 1c) encompassing three distinct motif categories. First, each design included at least one of the following motifs to create the near −180° turn necessitated by the design challenge: a large (>10 residue) bend such as a kink-turn (29), J5/5a from the P4-P6 domain (30), or an S-turn (31) (see, e.g., TWOWAY.1S72.20, TWOWAY.1GID.2 and TWOWAY.3BNQ.6 in Figure 1a; SI Table 2). Second, each design included at least one near-helical motif that was approximately, but not exactly, ‘straight’; an example is a set of three consecutive non-canonical base pairs (see, e.g., TWOWAY.1S72.51 in Figure 1a). Finally, some designs contained small motifs, such as a single adenine bulge or an A-A mismatch, used to make fine structural adjustments (see, e.g., TWOWAY.1S72.90 and TWOWAY.1S72.49 in Figure 1a). Previous work on RNA design, which was based on manual modeling by RNA experts, did not test these latter two categories of motifs, which are difficult to model without automatic tools, but appear to be necessary in natural RNAs for generating asymmetric structures in which small refinements of helical twists are required.

**Table 2:** Data collection, phasing and refinement statistics for miniTTR 6.

|  |  |
| --- | --- |
|  | Native |
| Space group | C2 |
| Cell dimensions |  |
| *a*, *b*, *c* (Å) | 233.372, 25.358, 42.861 |
| $\alpha$, $\beta$, $\gamma$ (°) | 90.0, 99.7, 90.0 |
| Wavelength | 1.58954 |
| Resolution (Å) | 2.55-115 (2.55-2.75) |
| *R*<sub>sym</sub> or *R*<sub>merge</sub> | 13.5(72.7) |
| *I* / σ*I* | 6.445(1.0) |
| Completeness (%) | 94.3(88.9) |
| Redundancy | 1.555(1.376) |
| **Refinement** |  |
| Resolution (Å) | 2.55-50.0 |
| No. reflections | 8467 |
| *R*<sub>work</sub> / *R*<sub>free</sub> | 22.9/27.66 |
| No. atoms | 3071 |
| RNA | 3049 |
| Ligand/ion | 10 |
| Water | 4 |
| *B*-factors |  |
| RNA | 40.915 |
| Ligand/ion | 33.284 |
| Water | 11.047 |
| R.m.s deviations |  |
| Bond lengths (Å) | 0.0077 |
| Bond angles (°) | 1.762 |

### Chemical Mapping of MiniTTRs to Probe TTR Formation

To probe the structures of the miniTTR designs generated by RNAMake, we first performed selective 2’-hydroxyl acylation analyzed by primer extension (SHAPE) and quantitative chemical mapping with dimethyl sulfate (DMS) (SI Figure 1) (32). For all 16 designs, we compared the SHAPE and DMS reactivity of each miniTTR RNA to its respective secondary structure. Of the 1386 nucleotides in the sixteen miniTTR constructs, 1367 (98.7%) were either reactive at target unpaired regions or protected at target helical residues, supporting the predicted secondary structures. All 19 outliers occurred at helix edges *(i.e*., flanking base pairs of motifs, SI Table 3). These data supported the formation of the expected secondary structures for all miniTTR designs (SI Figure 1).

To evaluate the formation of tertiary structure, we investigated the change in DMS reactivity of both tetraloop and tetraloop-receptor adenines as a function of Mg^2+^ concentration. Previous studies have demonstrated that TTR formation in the P4-P6 domain is strongly stabilized by Mg^2+^ (33–35). As a control for the unfolded state, the DMS reactivities of the tetraloop and tetraloop-receptor adenines of the TTR in the P4-P6 domain (A248, A151, A152 and A153) were measured and found to be 1.27, 0.72, 0.70, and 0.90, respectively, in 50 mM Na-HEPES, pH 8.0 buffer without Mg^2+^. Here and below, the values are normalized to the reactivity of the reference hairpin loops that flank each design (32). Upon the addition of 10 mM Mg^2+^, the adenines involved in the TTR became protected from DMS modification in the P4-P6 control (Figure 2d). As with this folding control, for 12 of the 16 designs (miniTTRs 1, 2, 5-7, 9-12 and 14-16), we observed a more than two-fold decrease in the reactivity of the TTR adenine residues (Figure 2d). These results were consistent with Mg^2+^-dependent TTR formation. The remaining designs (miniTTRs 3, 4, 8 and 13) did not demonstrate significant changes in DMS reactivity upon addition of 10 mM Mg^2+^, indicating that the TTR interaction did not form.

**Figure 2:**
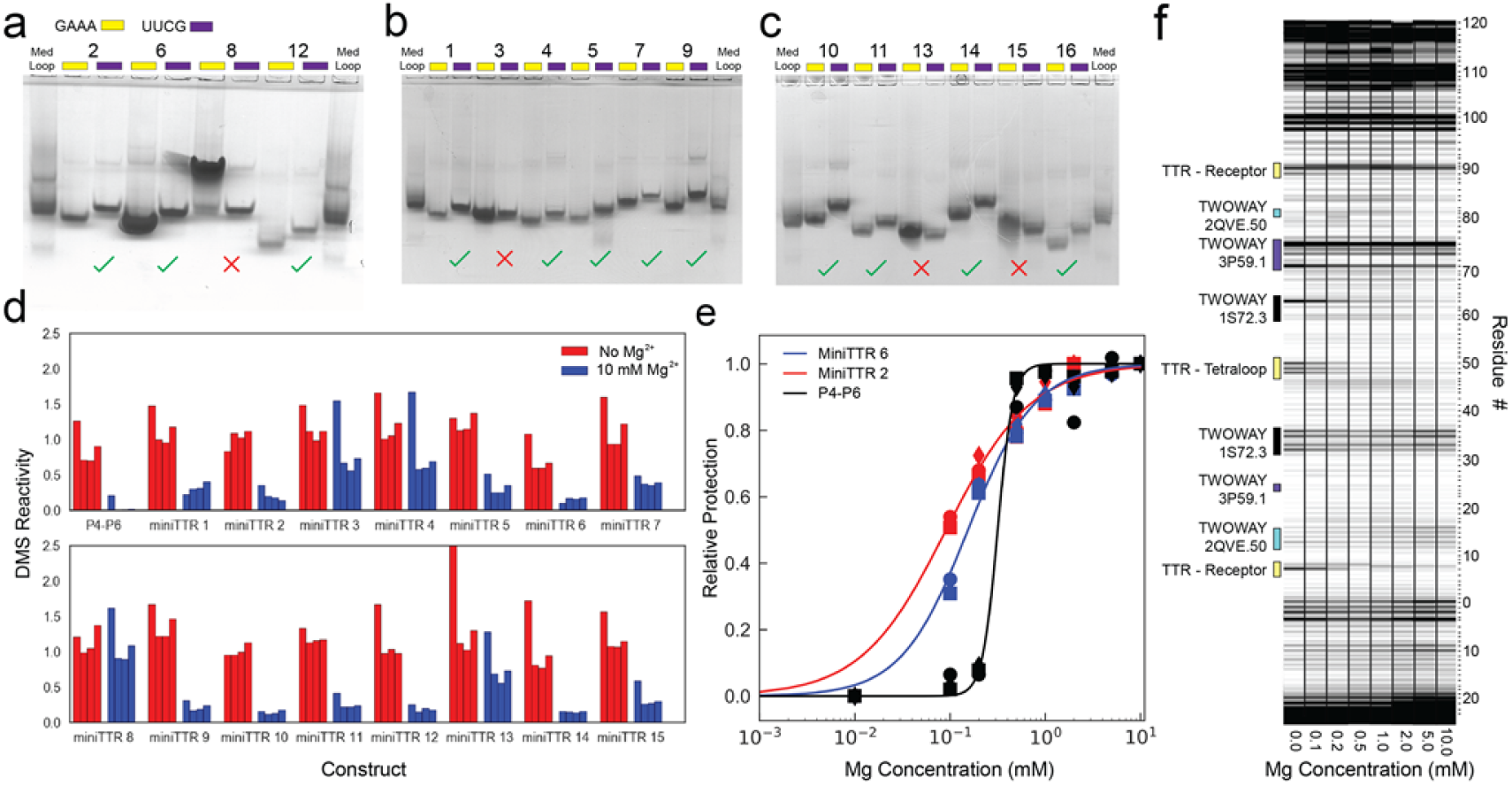
a-c) Native gel assay for each miniTTR design. Each design contains both a wild-type RNA with a GAAA tetraloop and a UUCG mutant that is unable to form the TTR interaction. d) Quantification of DMS reactivity with and without 10 mM Mg^2+^ for P4-P6 and all tested miniTTR constructs. e) Quantification of the change in DMS reactivity of adenosines involved in the TTR as a function of Mg^2+^ for miniTTR 6 (blue), miniTTR 2 (red) and P4-P6 (black). f) Raw data from the Mg^2+^ titration of miniTTR 2, highlighting the change in DMS reactivity in the TTR and the motifs used in the design.

To more precisely assess and compare folding stabilities, we determined how much Mg^2+^ was required to fold each of the miniTTRs that passed the screen described above (36). The miniTTR designs exhibited midpoints ranging from 0.12 to 7.0 mM Mg^2+^ (SI Table 4). The most stable designs were miniTTR 2 (Figure 2e and 2f) and miniTTR 6 (Figure 2e), with midpoints of 0.94 ± 0.39 and 0.12 ± 0.03 mM Mg^2+^ respectively. Their midpoints were similar to the Mg^2+^ midpoint measured for P4-P6 (0.38 ± 0.06 mM, Figure 2e). These observations suggested that at least two of the 16 automated designs achieved stabilities similar to that of the natural P4-P6 RNA fold, which is stabilized by the same tertiary contact, as well as by an additional tertiary contact (a metal core/metal core receptor) (28).

### Native Gel Mobility Shifts Due to TTR Formation

As an independent test of miniTTR folding, we replaced each RNA’s GAAA tetraloop with a UUCG tetraloop, which does not form the sequence-specific TTR tertiary contact (37) and is predicted to reduce the RNA’s mobility in non-denaturing polyacrylamide gel electrophoresis, as observed for the P4-P6 domain (38). Of the 16 miniTTR constructs tested, 12 designs displayed mobility shifts consistent with the formation of the TTR tertiary contact (Figure 2a-c, also see Table 1 for a comparison to chemical mapping). Constructs 4 and 15 exhibited mobility shifts that were inconsistent with our chemical mapping results. The UUCG mutant of miniTTR design 4 displayed a mobility shift, but it did not demonstrate a full two-fold decrease in TTR DMS reactivity, suggesting partial folding. Compared to its UUCG mutant, miniTTR design 15 in the wild-type form exhibited a wide, slow-mobility band, suggesting multimer formation. In all other cases, the electrophoretic mobility measurements were concordant with our quantitative SHAPE and DMS chemical mapping data, supporting the formation of the TTR and a compact tertiary fold.

### Small-angle X-ray Scattering Suggests Predominantly Stable Monomers

We carried out small-angle X-ray scattering (SAXS) measurements on our most stable designs, miniTTR 2 and miniTTR 6 (Figure 3). The observed scattering profiles of miniTTR 2 and miniTTR 6 agreed reasonably well with the profiles predicted from their corresponding RNAMake models, with χ^2^/d.o.f. values of 13 and 27, respectively (Figure 3a). These values are near values of 2–8 obtained from comparisons between predictions of RNA crystallographic models and scattering data (39–41), suggesting similar overall folds but with some local differences, an expectation confirmed for miniTT6 by X-ray crystallography (see next section).

**Figure 3:**
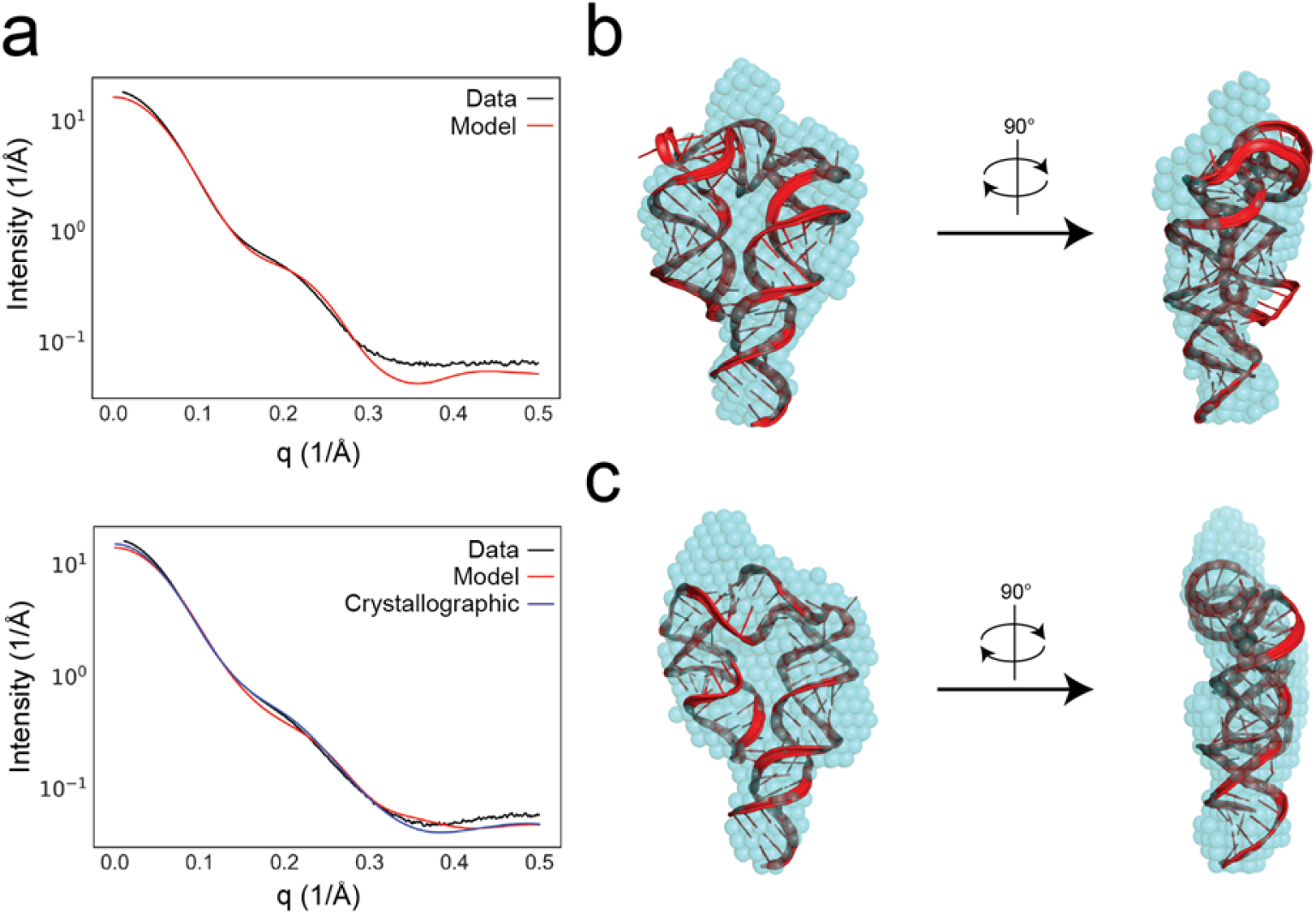
SAXS analysis of miniTTRs 2 and 6. a) Experimental intensity versus scattering vector for miniTTR 2 (top) and miniTTR 6 (bottom) compared to predictions based on models. b, c) Bead model derived from experimental scattering profiles overlaid with RNAMake models for miniTTR 2 (b) and miniTTR 6 (c).

To compare the coarse shapes of miniTTR designs 2 and 6 in solution to the predicted shapes, we generated low-resolution 3D representations of the experimental X-ray scattering profiles through the automated arrangement of ‘dummy atoms’ guided solely by the data (see Methods). For both constructs, the overall shapes of these unbiased shape reconstructions agreed well with the global folds of the RNAMake models (Figure 3b and 3c). Finally, size exclusion chromatography monitored by SAXS confirmed that <15% and <5% of miniTTRs 2 and 6, respectively, occur in higher-order structures (SI Figure 2). These results confirm that both the tested miniTTRs are primarily stable monomers, even at the RNA concentrations used for SAXS (>1 μM), again consistent with our RNAMake models.

### X-ray Crystallographic Structure Demonstrates Near-atomic Accuracy of Design

As the most stringent test of the RNAMake design algorithm, we sought to compare our modeled structures to atomic-resolution experimental structures. After targeting several of the miniTTR constructs for crystallization, we were able to obtain crystals of miniTTR 6 that diffracted at 2.55 Å resolution (I/σ of 1.0) (Table 2), and these data allowed for an atomic-level reconstruction. The SAXS profile predicted from these coordinates agreed with the SAXS data (χ^2^/d.o.f. = 9, Figure 3a), supporting the assumption that the crystal captured the solution conformation. The global fit between the crystallographic model and the structure predicted by RNAMake gave a heavy-atom RMSD of 4.2 Å. This value was somewhat less accurate than the heavy-atom RMSD of 3.4 ± 1.0 Å estimated from modeling the RNA’s thermal fluctuations (SI Figures 3 and 4), and the main source of deviation could be traced to a single feature: a triple mismatch (TWOWAY.1S72.62) (Figure 4e). The 3D model of this triple mismatch was derived from a ribosome crystal structure in which multiple bases flipped out from the motif to form A-minor interactions with flanking helices. In the context of the miniTTR, this motif did not flip out the bases; instead, it formed multiple consecutive non-canonical base pairs with high B-factors, presumably due to the absence of tertiary interaction partners. However, the other motifs of the design achieved near-atomic accuracy. Aligning the TTR from the model to the crystal structure (Figure 4b) gave a heavy-atom RMSD of 0.45 Å. Another motif selected by RNAMake in miniTTR 6 was TWOWAY.1S72.20, a kink-turn variant from the archaeal 50S ribosomal subunit (42). The RNAMake model agreed with the crystal structure with a heavy-atom RMSD of 2.0 Å. The overall topologies and interactions were nearly identical, and the flipped-out residues were also in the same orientation (Figure 4c). The last motif came from a viral internal ribosomal entry site domain incorporated into an RNA nanosquare (11). This motif exhibited a heavy-atom RMSD of 1. 28 Å between the RNAMake model and the crystal structure (Figure 4d). The only difference involved two unpaired residues that were stacked in the RNAMake model but flipped out in the miniTTR 6 crystal structure to form water-mediated contacts with other symmetry-related copies of the RNA in the crystal lattice. These changes were minor and did not affect the relative positioning or orientation of the helices connected by the motif (Figure 4d).

**Figure 4:**
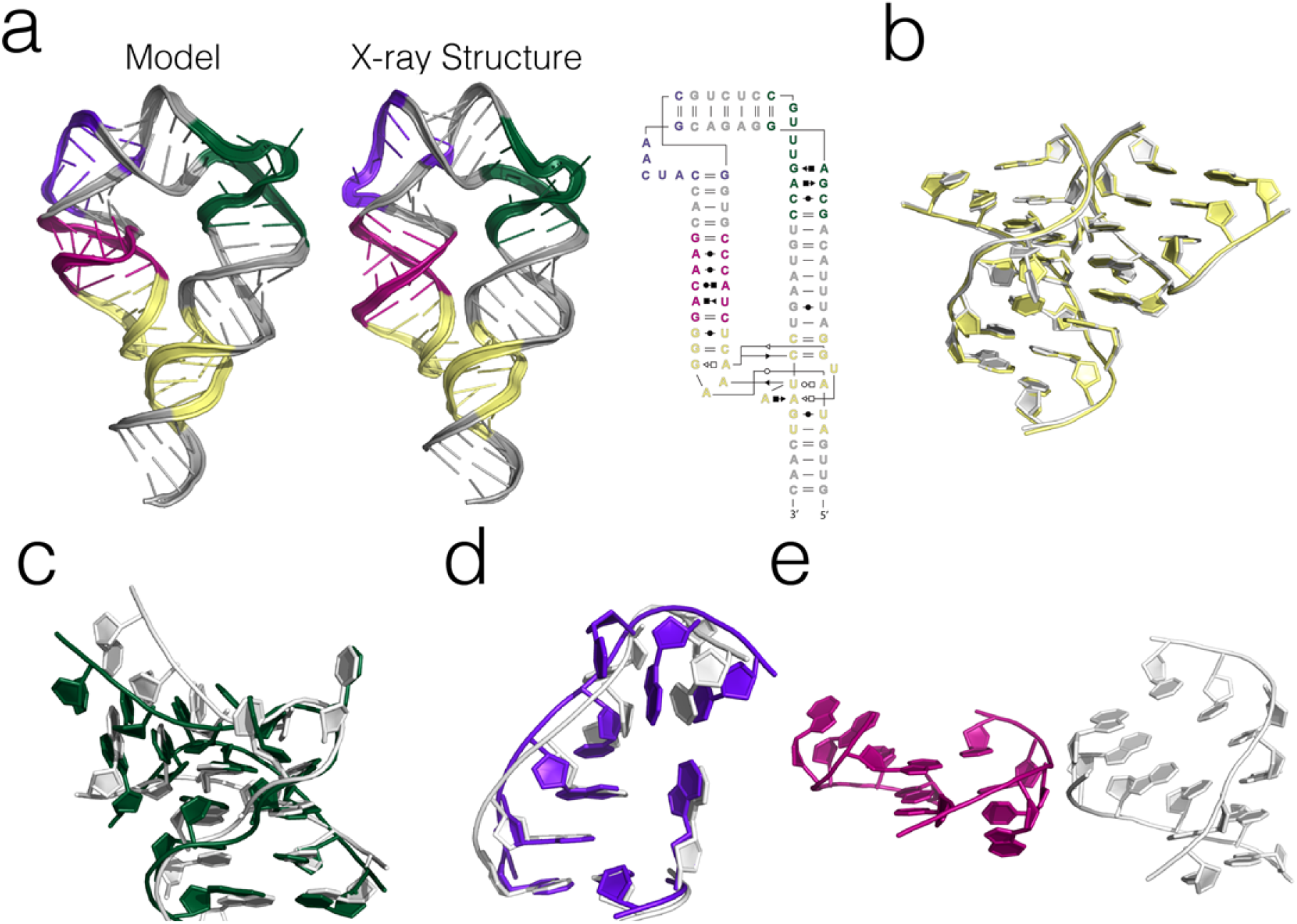
Comparison between the RNAMake model and X-ray crystallographic structure of MiniTTR 6. a) Overviews of both the entire RNAMake model, X-ray crystallographic and secondary structure of miniTTR 6. b) Overlay of the tetraloop/tetraloop-receptor (TTR) region. c) Overlay of the first motif selected by RNAMake. d) Overlay of the second motif selected by RNAMake. e) Comparison between the RNAMake model (left) and the X-ray crystal structure (right) of the last motif selected by RNAMake. In (b)-(e), the motif from the X-ray structure is in white, and the RNAMake model is in color.

### Tethered Ribosomes Designed Using RNAMake Support *Escherichia coli* Growth

After RNAMake’s success in the miniTTR structure design challenge, we sought to assess the algorithm’s ability to design molecules for function in a complex biological context. As a proof of concept, we designed RNA segments to tether the ribosome. The ribosome is a ribonucleoprotein machine dominated by two extensive RNA subunits. Previously we described the first successful construction of a ribosome with covalently tethered subunits that could support bacterial growth even in the absence of wild type ribosomes (23). In that work, several iterations of design were necessary to identify RNA tethers that support life in *E. coli* and were not cleaved by ribonucleases *in vivo*.

Here, we used RNAMake to computationally design RNA segments that tether the 50S and 30S ribosomal subunits together (SI Figure 5). The tethers were built between the H101 helix on the 50S subunit and the h44 helix on the 30S subunit (H101_h44_Tether), similar to the previous Ribo-T design (23). We designed nine unique constructs with RNAMake (Figure 5b) and successfully cloned seven. Each design utilizes between four and five motifs. The overlap between the motifs utilized in the miniTTRs and the H101_h44_Tether designs is low. Of the fourteen motifs used in the H101_h44_Tether designs, only three were also in the miniTTRs (TWOWAY.1NUV.3, TWOWAY.1S72.29 and TWOWAY.1S72.39). Nevertheless, like the miniTTRs, the H101_h44_Tether designs contained the same three distinct categories of motifs. First, all designs contained two large (>10 residue) bends (see, e.g., TWOWAY.1S72.29,

**Figure 5:**
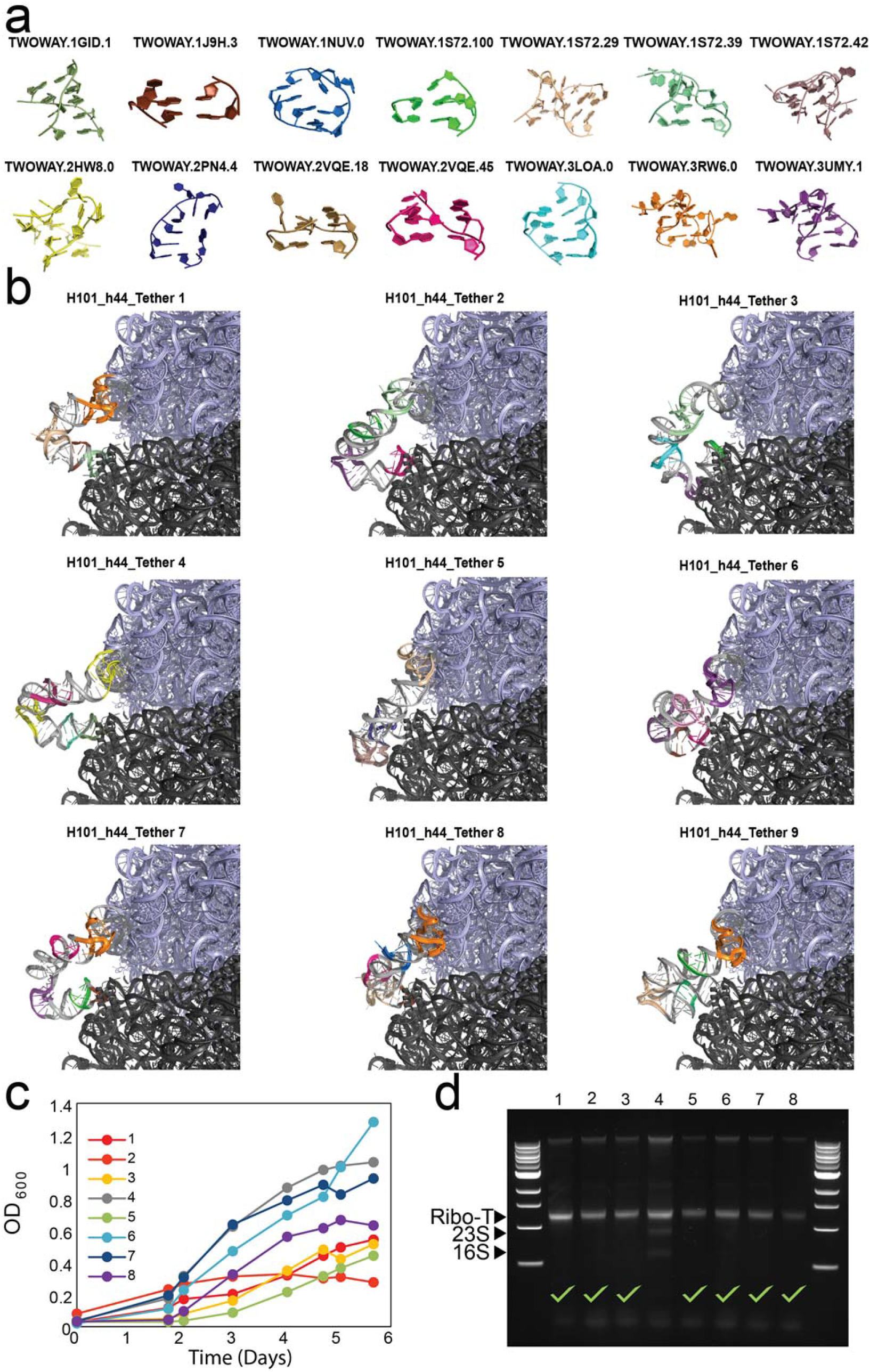
Summary of the RNAMake-based design of a functional tether between the 50S and 30S ribosomal subunits. a) All unique motifs included in the H101_h44_Tether designs. b) Ensemble of all RNAMake H101_h44_Tether designs. Each is colored according to the motifs (panel (a)) that were used to generate it. c) Culture density of *E. coli* cells containing RNAMake H101_h44_Tether design 4. d) Gel assay of 8 distinct colonies, verifying that all but clone 4 have a Ribo-T rRNA-sized band with loss of wild-type 23S and 16S rRNA bands.

TWOWAY.1S72.39 and TWOWAY.1S72.42 in Figure 5a). This characteristic differed from that of the miniTTR designs, where it was possible to only have one bend. Furthermore, unlike the miniTTRs, each design contained at least one near-helical motif (see, e.g., TWOWAY.1NUV.3, TWOWAY.2VQE.46, TWOWAY.3LOA.0 in Figure 5a) and one small motif (see, e.g., TWOWAY.1S72.100, TWOWAY.1J9H.3 in Figure 5a).

To test RNAMake’s accuracy in designing functional RNA, each H101_h44_Tether design (Figure 5a) was cloned into the pRibo-T plasmid (23) and used to replace the wild-type ribosomal rRNA plasmid in the SQ171fg strain (23). After 48 hours at 37 °C, no colonies were visible on the plates. Fresh plates were replica plated and incubated for a further 72 hours at 37 °C, after which colonies appeared on the H101_h44_Tether design plate. Eight colonies were picked and checked for the loss of the wild-type rRNA plasmid. Growth curves were generated in liquid culture at 37 °C (Figure 5c). After 5 days of growth, total RNA was extracted and analyzed by gel electrophoresis. All clones (except clone 4) showed a clean Ribo-T-sized band and no detectable wild-type 23S and 16S rRNA bands (Figure 5d), indicating the formation of stable tethered ribosomes as previously demonstrated (23). Plasmids were extracted from these colonies, and their sequences were each confirmed as the correct H101_h44_Tether design 4 plasmid. The growth curves of the 7 successful clones (excluding clone 4) exhibited doubling times of 1. 5 ± 0.8 days and a maximum OD_600_ of 0.7 ± 0.3. While slower than wild type *E. coli*, the first successful version of Ribo-T was also slower growing before mutational optimization (23). Taken together, these data demonstrate that tethered ribosomes with automatically designed tethers using RNAMake can fully support *E. coli* growth in the absence of wild-type ribosomes.

## Discussion

### Automated 3D Design with a High Success Rate

We have demonstrated that RNAMake can automate the design of new RNA structures, requiring only the start and end points of the desired structure. Taking a classic and common problem in RNA 3D design, the translational and rotational alignment of partner segments to form a target tertiary contact, RNAMake generated 11 out of 16 successful designs that passed our chemical mapping and native gel mobility tests. Furthermore, miniTTR 2 and 6 achieved stability at low [Mg^2+^], similar to or better than the natural TTR-containing P4-P6, despite using fewer nucleotides (97 and 94 nts respectively, compared to the 157 nts in the P4-P6 domain) and containing fewer tertiary contacts (the P4-P6 domain contains a metal-core/metal-core receptor in addition to a TTR). One possible reason why these two miniTTR designs were more stable than the P4-P6 domain at lower [Mg^2+^] is that the motifs in the miniTTR designs were chosen rationally to achieve a given orientation, while the motifs in the P4-P6 domain were naturally selected to support the broader folding landscape and multi-step functional cycle of the larger *Tetrahymena* self-splicing group I intron. Interestingly, miniTTRs have shallower Mg^2+^ dependence than the P4-P6 RNA (Figure 2); designing sharper Mg^2+^ dependences, which are characteristic of many natural tertiary RNA structures, will be an important future challenge. Our results suggest that it is possible to computationally generate RNAs that are just as or more stable than naturally occurring RNAs, analogous to results in protein design (43, 44).

### Expanding the Experimentally Vetted ‘Building Blocks’ of RNA 3D Design

In RNA nanotechnology efforts to date, utilizing a previously untested motif in 3D design has required trial-and-error testing to verify that the RNA remains in a well-defined 3D structure in new contexts. Due to the amount of human and experimental effort required, the inclusion of novel motifs has been slow, with researchers generally drawing from a limited set of motifs previously verified to be modular and reliable, and these motifs are typically deployed one at a time within highly repetitive arrangements rather than being combined into complex asymmetric combinations. This process has generated simple RNAs that lack the complexity of natural RNA machines.

In our miniTTR study, automated computational design resulted in RNAs containing a total of 23 motifs, 20 of which had not been used previously in the 3D rational design of RNA. The motifs ranged from a single bulged A to complex motifs such as kink-turn variants and S-turns (Figure 1a, SI Table 2). Of these new motifs, a large majority (18) appeared in designs validated by our biochemical experiments, but two others gave informative discrepancies. One motif, an A-A mismatch (TWOWAY.1S72.49, Figure 1a), appeared in all four miniTTR designs that did not form TTRs (3, 4, 8, 13). Unlike other A-A mismatches that form a non-canonical base pair (45, 46), the adenines in the TWOWAY.1S72.49 structure are not paired and may reflect a high-energy structure that should not be used to design stable assemblies. The other motif with an interesting structural deviation was a triple mismatch (TWOWAY.1S72.62, Figure 1a). Chemical mapping suggested that this motif retained its anticipated conformation at nucleotide resolution in miniTTR 6, but crystallographic analysis showed the motif’s structure to be incorrect at atomic resolution (Figure 4e). TWOWAY.1S72.62’s alternative conformation is likely due to the lack of the tertiary contacts provided by its parent ribosomal context but not the miniTTR design. Interestingly, this structural change was still compatible with the global folding of the miniTTR 6 design, which, in fact, was our most stable design, as assessed by Mg^2+^ titration. This result suggests that residual uncertainties in RNAMake’s motif library will not preclude the consistent design of asymmetric structures at nucleotide resolution. Our experimental tests of the remaining 18 modular motifs, including two verified at atomic resolution, expand both the number and diversity of motifs available for RNA nanotechnology.

### RNAMake Accelerates the Design of *in vivo* RNA Nano-machines

The generation of a hybrid ribosomal RNA containing the sequences of both the small and large rRNA subunit sequences linked together by an engineered two-stranded tether significantly expanded the promise of synthetic biology. Yet, without the assistance of 3D modeling software such as RNAMake, the time invested in its development was substantial, requiring over a year of experimenting with different tether sequences (25). Utilizing RNAMake, we greatly simplified the design process. In our first attempt, we generated nine unique tether designs and successfully cloned seven. Of these, one design was able to support *E. coli* growth in the absence of wild-type ribosomes, while remaining uncut *in vivo*. Although its growth rate was slow (doubling every 1.5 ± 0.8 days), achieving success *in vivo* on the first attempt suggests that RNAMake can significantly accelerate design of RNA machines. As in the case of the first Ribo-T, we can now use both rational design and selection techniques to further optimize this tether for the multiple states in the ribosome’s functional cycle. RNAMake enables the computational design of diverse molecules to solve new problems, raising the prospect of increasing RNA design complexity beyond what has been achieved with manual methods.

## Acknowledgments

We thank S. Bonilla for assistance in performing native gel assays and A. Watkins for discussions about ribosome tether design. This work was supported by National Institutes of Health grant R01 GM100953 (to R.D.), Ruth L. Kirschstein National Research Service Award Postdoctoral Fellowships GM112294 (to J.D.Y.) and GM100953 (to D.E.), National Institute of General Medical Sciences grant R35 GM118070 (to J.S.K.), Army Research Office W911NF-16-1-0372 (to M.C.J.), National Science Foundation grant MCB-1716766 (to M.C.J.), the David and Lucile Packard Foundation (to M.C.J.) and National Institutes of Health grant P01 GM066275 (to D.H.).

## Contributions

R.D. and J.D.Y. conceived of the study. J.D.Y. developed RNAMake and generated the models and sequences used throughout the study. A.N.O. and W.K. performed the experimental chemical mapping, titrations and native gel assays. J.D.Y. analyzed and processed the results from these experiments. X.S. performed the small-angle X-ray scattering on miniTTRs 2 and 6. D.E. solved the crystal structure of miniTTR 6 and was assisted by D.C. and J.S.K. in preparing the RNA and in the analysis. E.D.C. and M.C.J. made and tested the RNAMake-designed Ribo-T tethers. J.D.Y. and R.D. wrote the paper, with participation by all authors.

## Methods

### Software Availability

All software and source code used in this work are freely available for non-commercial use. RNAMake software and documentation are at <u>https://github.com/jyesselm/RNAMake</u>. EteRNAbot is available at <u>https://github.com/EteRNAgame/EteRNABot</u>.

### Building the Motif Library

To build a curated motif library of all RNA structural components, we obtained the set of non-redundant RNA crystal structures managed by the Leontis and Zirbel groups (18) (version 1.45: http://rna.bgsu.edu/rna3dhub/nrlist/release/1.45(weblink)). This set specifically removes redundant RNA structures that are identical to previously solved structures, such as ribosomes crystallized with different antibiotics. We processed each RNA structure to extract every motif with Dissecting the Spatial Structure of RNA (DSSR) (47) with the following command:

~~~
x3dna-dssr -i file.pdb -o file_dssr.out
~~~

We manually checked each extracted motif to confirm that it was the correct type, as DSSR sometimes classifies tertiary contacts as higher-order junctions and vice-versa. For each motif collected from DSSR, we ran the X3DNA **find_pair** and **analyze** programs to determine the reference frame for the first and last base pair of each motif to allow for alignment between motifs:

~~~
find_pair file.pdb 2> /dev/null stdout | analyze stdin >& /dev/null
~~~

In addition to the motifs derived from the PDB, we also utilized the make-na web server (<u>http://structure.usc.edu/make-na/server.html</u>) to generate idealized helices of between 2 and 22 base pairs in length (26). All motifs in these generated libraries are bundled with RNAMake and are grouped together by type (junctions, hairpins, etc.) in sqlite3 databases in the directory RNAMake/rnamake/resources/motif_libraries/(weblink).

### Automatically Building New RNA Segments

RNAMake seeks a path for RNA helices and noncanonical motifs that can connect two base pairs separated by a target translation and rotation. We developed a depth-first search algorithm to discover such RNA paths. The algorithm is guided by a heuristic cost function *f* inspired by prior manual design efforts (5, 11) and is composed of two terms:

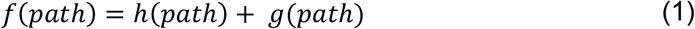

The first term, *h(path)*, describes how close the last base pair in the path is to the target base pair; *h(path)* = 0 corresponds to a perfect overlap in translation and rotation. The functional form for *h(path)* depends on the spatial position of each base pair’s centroid *d* and an orthonormal coordinate frame *R* defining the rotational orientation of each base pair (48):

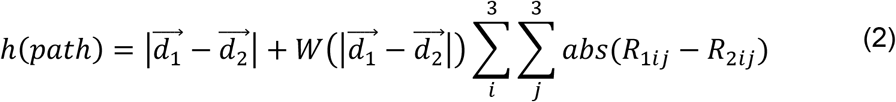

Here, *W(d)* is:

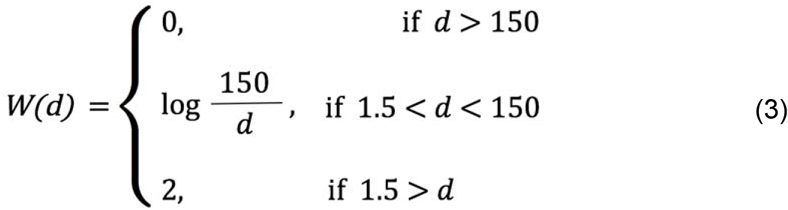

The weight *W(d)* reduces the importance of the current base pair and the target base pair with similar alignment when they are spatially far apart. This term conveys the intuition that aligning the two coordinate frames becomes important only as the path of the motif and helices approaches the target base pair. RNAMake readily allows for the exploration of alternative forms of the cost function terms in (2) and (3), including more standard rotationally invariant metrics to define rotation matrix differences (49) or base-pair-to-base-pair RMSDs based on quaternions (50), but these were not tested in the current study.

The second term in the cost function (1) is *g(path)*, which parameterizes the properties of the non-canonical RNA motifs and helices comprising the path at each stage of the calculation

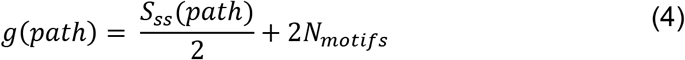

where *S_ss_* is a secondary structure score for all the motifs and helices in the path. This SS favors longer helices as well as motifs with frequently recurring base pairs, as follows. All base pairs found in the RNA motif are scored based on their relative occurrences in all high-resolution crystal structures; all unpaired residues receive a penalty, and Watson-Crick base pairs receive an additional bonus score (values are given in SI Table 5). In addition to the secondary structure score, *N_motifs_* penalizes the total number of motifs in the path, here taken as the number of non-canonical motifs plus the number of helices (independent of helix length).

The search adds motifs and helices to the path in a depth-first manner, while the total cost function *f(path)* decreases, back-tracking if *f(path.)* increases. Any solutions with *h(path)* less than 5, i.e., overlap at approximately nucleotide resolution between the path’s last base pair and the target base pair, are accepted into a list of final designs.

The balance between *g(path.)* and *h(path.)* allows RNAMake to reduce the number of motif combinations considered, finding most solutions in a few seconds. For each solution, we then used EteRNAbot, a secondary structure optimization algorithm that has undergone extensive empirical tests (27), to fill in helix sequences.

An example RNAMake command line is the following:

~~~
design_rna -pdb p4p6_short.pdb -start_bp A146-A157 -end_bp
A221-A252-designs 1000000
~~~

Here, `p4p6_short.pdb` is P4-P6 with residues 233 to 241 removed. This command runs the RNAMake design algorithm to build a new RNA segment between the base pair consisting of nucleotides 146 and 157 and the base pair consisting of nucleotides 221 and 252, also on chain A. The `design_rna` application automatically removes the nucleotides between these two ends, leaving only the two segments of the TTR remaining.

### RNA Structural Probing and miniTTR Data Analysis

DNA oligonucleotides were designed with Primerize (51), ordered from IDT (Integrated DNA Technologies) and used to generate double-stranded DNA templates using PCR assembly. DNA template and RNA transcript preparation and quality checks were carried out as previously described (32, 52).

Chemical mapping (DMS and SHAPE) was performed as previously described (32, 52). Briefly, modification reactions were performed in a 20 μL volume containing 1.2 pmol of RNA and 50 mM Na-HEPES (pH 8.0). Before the chemical modifier was added, the RNA was heated to 90 °C for 3 minutes, then left on the bench top to cool to room temperature and then folded for 20 min in 10 mM MgCl_2_ and 50 mM Na-HEPES (pH 8.0). To chemically modify the RNA, either 5 μL of DMS (1% v/v in 10% ethanol) or 1M7 (5 mg/mL in anhydrous DMSO) were added to each reaction to a total volume of 25 μL. After 5 minutes of incubation at room temperature, the reactions were quenched with 0.5 M Na-MES (pH 6.0). After quenching, poly(dT) magnetic beads (Ambion) and FAM-labeled Tail2-A20 primers were added for reverse transcription. Samples were separated and purified using magnetic stands, washed twice with 70% ethanol, and air-dried. The beads were resuspended in ddH_2_O and reverse transcription mix, then incubated at 48 °C for 30 min. RNA was degraded by adding 1 volume of 0.4 M NaOH and incubating at 90 °C for 3 minutes; it was then cooled and neutralized with an additional volume of acid quench (prepared as 2 volumes of 5 M NaCl, 2 volumes of 2 M HCl, and 3 M sodium acetate, pH 5.2). Fluorescently labeled cDNA was recovered by magnetic bead separation, rinsed twice with 70% ethanol and air-dried. The beads were resuspended in Hi-Di formamide containing ROX-350 ladder (Applied Biosystems), then loaded on a capillary electrophoresis sequencer (ABI3130, Applied Biosystems).

The HiTRACE 2.0 package was used to analyze the CE data, available as a MATLAB toolbox at <u>https://github.com/hitrace</u> (53) and a web server at <u>http://hitrace.org</u> (54). Electrophoretic traces were aligned and baseline subtracted using linear and nonlinear alignment routines as previously described (53). Reactivities were determined by fitting these traces to sums of Gaussian peaks, followed by background subtraction, signal attenuation correction, and normalization to flanking reference hairpins (32).

To estimate Mg^2+^ titration midpoints, the relative protection values (f_j_^i^) for each residue *j* in the TTR at each Mg^2+^ concentration *i* were calculated. The quantitative DMS reactivity of the folded and unfolded state of each TTR residue was taken from P4-P6.

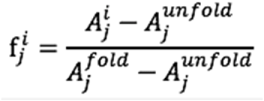

These values were then fit to the Hill equation:

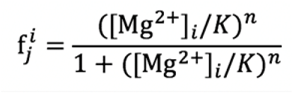

For each data set, global Hill fits were generated using scipy’s curve_fit function, and errors were estimated through bootstrapping.

### Native Gel Shift Assays

Native gel shift assays were conducted using a BioRad Criterion™ Cell gel cassette. Gels were cast using 11% acrylamide and 1X THEM buffer (pH 7.2) (10 mM MgCl_2_) and polymerized by adding 10% ammonium persulfate (300 μL) and TEMED (30 μL) to 30 mL of gel mix. Following polymerization, the gel apparatus was set up in a 4 °C cold room and fully immersed in an ice bath until the gel and buffer apparatus were cooled to approximately 4 °C. Then, RNA constructs were prepared for folding by incubating them in folding buffer consisting of 100 mM Na-HEPES (pH 7.5), 10 mM MgCl_2_, and 0.5 mM spermidine. 100 μg of RNA was mixed with folding buffer in the absence of spermidine and incubated at 65 °C for 3 min. The solution was cooled at room temperature for 10 min, and then 0.5 mM spermidine was added. The RNAs were then vortexed for 10 seconds and centrifuged at 4,000 rpm for 10 min. Immediately following folding, loading dye was added to the RNA solutions, and the samples were directly loaded onto the cooled gel apparatus. The gel was run at 15 watts for 1.5 hours, and the temperature of the gel apparatus was strictly monitored to avoid overheating. After electrophoresis, the gel was removed from the cassette, carefully placed in a glass pan and incubated with 150 mL of Stains-All on an orbital shaker for 15 min. For de-staining, the Stains-All was removed, and the gel was rinsed with deionized water and subsequently incubated in fresh deionized water on an orbital shaker for 20 min; the gel was then immediately imaged on a document scanner.

### SAXS Measurements, Analysis and Modeling

RNA transcripts were purified using an Agilent 1260 Infinity HPLC using a gradient of 13–23% buffer B (100 mM TEAA (pH 7.0) and 50% acetonitrile) in buffer A (100 mM TEAA (pH 7.0) and 0.2% sodium azide) over a Varian PLRP-S 1000 Å 8 μm 150 × 7.5 mm column. Fractions containing miniTTR constructs were pooled, concentrated, and buffer-exchanged three times into water using Amicon Ultra concentrators (Millipore). The RNA was then quantified and stored at -20 °C until use.

Small-angle X-ray scattering measurements were carried out at Bio-SAXS beamline BL4-2 at the Stanford Synchrotron Radiation Lightsource (SSRL). Scattering data were collected with a 1.7 m sample-to-detector distance and a beam energy of 11 keV (wavelength of 1.127 Å). RNA samples were first buffer-exchanged into running measurement buffer solution consisting of 70 mM Tris-HCl (pH 7.4), 160 mM NaCl, 10 mM MgCl_2_ and 5 mM DTT using Amicon Ultra centrifugal filters (10K cutoff, Millipore). Approximately 50 μL of the buffer-exchanged RNA (5 mg/mL) was then loaded onto a 24 mL Superdex 200 size-exclusion column (GE Healthcare) that had been pre-equilibrated with the running measurement buffer solution, then run at a flow rate of 0.05 mL/min using an AKTA Ettan FPLC (GE Healthcare). The elution was directed to the sample flow path for immediate SAXS data collection every 5 seconds, with a 2 second exposure time. The SAXS images were processed using the SASTOOL program. The first 100 images were used to create the buffer scattering profiles. The segment of the main elution peak with constant, scale-independent scattering profiles was used to calculate the sample scattering profiles.

The SAXS profiles of the miniTTR 2 and 6 RNAMake models were predicted and compared with the experimental profiles (Figure 3a) using FoXS (55). 3D bead models of miniTTR 2 and 6 were generated using DAMMIF and DAMMIN (56, 57) and overlaid with their corresponding RNAMake models in PyMOL.

### Materials and Methods for X-ray Crystallography

RNA used for crystallization was transcribed with T7 RNA polymerase from PCR-generated double-stranded DNA templates as described in (58). These templates were ordered from IDT as gBlocks with, in the 5′ to 3′ direction, a T7 promoter sequence, hammerhead ribozyme, miniTTR sequence of interest, and HDV ribozyme. RNA transcripts were ethanol precipitated overnight, washed with 70% ethanol and dissolved in water. RNA transcripts were purified from ribozymes and uncleaved products using PAGE purification. RNAs were eluted overnight at 4 °C, concentrated, and buffer-exchanged three times into water using Amicon Ultra concentrators (10K cutoff, Millipore). RNA was quantified and then stored at −20 °C until use.

### MiniTTR Crystallization

Purified miniTTR 6 RNA diluted in buffer A (30 mM HEPES (pH 7.5), 20 mM MgCl2, and 100 mM KCl) was incubated at 65 °C for 2 min, centrifuged at 13,000 rpm for 2 min, and snap-cooled on ice for approximately 5 min before moving to 25 °C to set up crystallization trays. Within 2-4 weeks, miniTTR 6 crystallized at 25 °C as plates or clusters of plates via sitting-drop vapor diffusion by mixing 2 μL of miniTTR 6 at a concentration of 100 μM with 3 μL of crystallization solution containing 40 mM sodium cacodylate (pH 5.5), 20 mM MgCl_2_, 2 mM cobalt hexammine, and 40% 2-methyl-2,4-pentanediol (MPD). Crystals of miniTTR 6 grew to maximum dimensions of 700 x 700 x 20 μm and were stabilized and cryogenically protected by increasing the MPD to a final concentration of 44%. Crystals were flash-frozen by plunging into liquid nitrogen.

Diffraction data were collected at 100 K using synchrotron X-ray radiation at beamline 4.2.2 of the Advanced Light Source, Lawrence Berkeley National Laboratory (Berkeley, CA). The data were processed and scaled using X-ray Detector Software (XDS) (59). The scaled data were handled using Collaborative Computational Project programs (60).

### Structure Determination and Refinement

The initial structural determination of the miniTTR in the C2 space group was carried out from molecular replacement (MR) in Phaser (CCP4) searching for one copy of a 31-nucleotide model of only the tetraloop and receptor with the identical sequence (60). The rotational and translational Z-scores were somewhat low, 4.6 and 5.9 respectively, but the maps were of sufficient quality to enable the iterative building of all the residues into the 2F_o_-F_c_ and F_o_-F_c_ maps. Composite omit maps in PHENIX were used to help confirm the model and reduce model bias from the initial MR solution (61). The models were built using COOT (62) and refined using REFMAC5 and PHENIX (60). The final model was refined in REFMAC5 (60), and the overall R_work_ and R_free_ were refined to 22.9% and 27.7%, respectively. The structure derived from the miniTTR was refined to 2.55 Å against a data set scaled to an overall I/σ of 1.0 at the highest resolution shell with 98.5% completeness. Final crystallographic statistics can be found in Table 2. The crystal structures of miniTTR 6 have been deposited in PDB, ID 5VOQ. All structural figures were prepared using PyMOL (<u>http://www.pymol.org/</u>).

### Design and Cloning of Novel Tethers for a Ribosome with Tethered Subunits

The designed tethers (SI Table 6) were cloned into plasmid pRibo-T-A2058G (23) using the primers in SI Table 7. The backbone was generated for each design using forward (f) and reverse (r) primer pairs (noted with “bb”) in SI Table 7 in separate PCR reactions using plasmid pRibo-T as a template (23), Phusion polymerase (NEB), and 3% DMSO. PCR cycling was as follows: 98 °C for 3 min; 25 cycles of 98 °C for 30 sec, 55 °C for 30 sec, 72 °C for 2 min; and 72 °C for 10 min. Circularly permuted 23S ribosomal RNA (rRNA) was generated with forward and reverse primer pairs (noted with “23S” in SI Table 7), the pRibo-T template, and the same PCR conditions as described above. Each PCR reaction was purified by gel extraction from a 0.7% agarose gel with an E.Z.N.A. gel extraction kit (Omega). Each purifed backbone (50 ng) was assembled with the respective 23S insert in 3-fold molar excess using Gibson assembly (63). Assembly reactions were transformed into POP2136 cells, and the cells were grown at 30 °C overnight. Colonies were picked and plasmids were isolated using an E.Z.N.A. miniprep kit (Omega) and confirmed with full plasmid sequencing by ACGT, Inc.

### Replacement of Wild-type Ribosomes with RNAMake Ribo-T

Each purified plasmid (1 μL) was separately transformed into electrocompetent SQ171fg cells containing pCSacB (23). Cells were recovered in 1 mL of SOC media at 37 °C with shaking for 1 hour. Fresh SOC (1.85 mL) supplemented with 50 μg/mL carbenicillin and 0. 25% sucrose was inoculated with 250 μL of recovered cells and incubated overnight at 37 °C with shaking. Cultures (10% and 90%) were plated on LB agar plates supplemented with 50 μg/mL carbenicillin, 5% sucrose and 1 mg/mL erythromycin and incubated at 37 °C.

After 48 hours with no visible colonies, the plates were replica plated onto fresh LB agar plates supplemented with 50 μg/mL carbenicillin, 5% sucrose and 1 mg/mL erythromycin and incubated at 37 °C. After 72 additional hours, colonies appeared on the plate containing H101_h44_Tether design 4. Eight colonies were streaked onto LB agar supplemented with 50 μg/mL carbenicillin and 1 mg/mL erythromycin and LB agar supplemented with 30 μg/mL kanamycin (to confirm loss of the pCSacB plasmid) and were also used to inoculate 5 mL of LB supplemented with 50 μg/mL carbenicillin and 1 mg/mL erythromycin. Plates were incubated at 37 °C, and cultures were incubated at 37 °C with shaking. The OD_600_ of the cultures was tracked to generate growth curves (Biochrom Libra S4 spectrophotometer). After 5 days at 37 °C, total RNA was extracted using an RNA extraction kit from Qiagen. Total RNA was analyzed by gel electrophoresis on a 1% agarose gel with GelRed. Total plasmid was extracted from saturated 5 mL cultures with an E.Z.N.A. miniprep kit (Omega) and sequenced to confirm the correct H101_h44_Tether design 4 sequence.

